# Atrophy associated with tau pathology precedes overt cell death in a mouse model of progressive tauopathy

**DOI:** 10.1101/2020.06.05.136663

**Authors:** Christine W. Fung, Jia Guo, Helen Y. Figueroa, Elisa E. Konofagou, Karen E. Duff

## Abstract

In the early stages of Alzheimer’s disease (AD), tau pathology first develops in the entorhinal cortex (EC), then spreads to the hippocampus and at later stages, to the neocortex. Pathology in the neocortex correlates with impaired cognitive performance. Overall, tau pathology correlates well with neurodegeneration but the spatial and temporal association between tau pathology and overt volume loss is unclear. Using *in vivo* magnetic resonance imaging (MRI) with tensor-based morphometry (TBM) we mapped the spatio-temporal pattern of structural changes in a mouse model of AD-like progressive tauopathy. A novel, co-registered *in vivo* MRI atlas identified particular regions in the medial temporal lobe (MTL) that had significant volume reduction. The medial entorhinal cortex (MEC) and pre-/para-subiculum (PPS) had the most significant atrophy at the early stage, but atrophy then spread into the hippocampus, most notably, the CA1, dentate gyrus (DG) and subiculum (Sub). TBM-related atrophy in the DG and Sub preceded overt cell loss that has been reported in *ex vivo* studies in the same mouse model. By unifying an *ex vivo* 3D reconstruction of tau pathology with the TBM-MRI results we mapped the progression of atrophy in the MTL with the corresponding spread of tau pathology. Our study shows that there is an association between the spread of tau pathology and TBM-related atrophy from the EC to the hippocampus, but atrophy in the DG and Sub preceded overt cell loss.

**One Sentence Summary:** Spread of tau pathology in a mouse model of Alzheimer’s disease assessed by MRI was associated with reduced brain tissue volume but not neuronal loss.

## Introduction

One of the major pathological hallmarks of Alzheimer’s disease (AD) and the primary tauopathies that cause frontotemporal degeneration (FTD-tau) is the accumulation of insoluble, hyperphosphorylated tau protein into intraneuronal neurofibrillary tangles (NTFs) (*1*), threads, and inclusions. In healthy neurons, the tau protein stabilizes axonal microtubules that are necessary for proper neuronal function. In AD and the tauopathies, the abnormal accumulation of tau protein in somatodendritic compartments is associated with axonal and synaptic dysfunction, cerebral atrophy, neuronal loss, and ultimately clinical and functional decline. In the earliest stages of AD, tauopathy begins in the entorhinal cortex (EC). Tau pathology then develops in the hippocampus, and the neocortical areas at later stages, which correlates with cognitive impairment (*2*). However, it is not yet fully understood how – and to what extent – the progressive spread of tau pathology is associated with axonal and synaptic dysfunction, and its relationship with cerebral atrophy and neuronal loss over time.

Imaging techniques such as positron emission topography (PET) and magnetic resonance imaging (MRI) have been used to explore the relationship between the distribution of tau pathology and atrophy respectively, in human AD (*3-11*). However, tau PET ligands have only recently been applied in human AD and longitudinal studies are limited. As human AD is characterized by the accumulation of amyloid plaques and tau neurofibrillary tangles (as well as synuclein and TDP43 inclusions in many cases of AD), the impact of tau pathology per se on degeneration is unclear. To address this, transgenic tau mice were created to investigate the spatial and temporal spread of tauopathy and its role in pathogenesis (*12-14*). Previous studies have shown that tau pathology originating in the EC can spread transynaptically from regions of primary vulnerability to secondary affected areas along neuroanatomically connected routes, and the idea that tauopathy spreads through the brain has gained credibility. However, it has not been demonstrated that pathology spread is accompanied by neurodegeneration in secondary affected regions, and the temporal and spatial relationship between pathology and neurodegeneration in models of progressive spread of tauopathy has yet to be shown.

Our aim was to elucidate the relationship between the pathological spread of tau and the associated spread of atrophy (volume reduction). To do this, we used EC-Tau mice at 20-24 months old (where the tauopathy represents an early stage in human AD tauopathy) and 30-36 months old (which represents a moderate stage). We utilized tensor-based-morphometry (TBM) with MRI to observe the global and regional structural changes between the EC-Tau mice and their age-matched controls. TBM is an imaging technique used to localize regions of shape differences based on nonlinear deformation fields that align, or warp, images to a common anatomical template (*15*). The Jacobian map is the determinant of the Jacobian matrix of a deformation field and quantifies the local change in volume at the voxel level in order to evaluate whether volume loss (or growth) has occurred. In this study, we not only show that the atrophy spreads from the EC to the hippocampus alongside tau pathology, but we also identify areas in the hippocampus with atrophy at the later stage that showed no overt neuronal loss in *ex vivo* studies. In addition, we visualized this progression of atrophy and the spread of tau pathology in 3D thus demonstrating that the spread of tau pathology is associated with significant TBM-related reductions in volume, but precedes overt cell loss at the moderate stage.

## Results

### Significant volume reduction in MTL of EC-Tau mice

Voxel-based analyses were performed to determine the amount of significant volume reduction (atrophy) in the EC-Tau group compared to their age-matched control group. A three-dimensional rendering of the mouse brain was generated using 3D Slicer software. As previously described (*16*), an *in vivo* MRI mouse atlas was utilized to locate different sub-regions in the mouse brain, specifically the cortex and hippocampus. In our analyses, we focused on significant atrophy in the medial temporal lobe (MTL). The MTL consists of the entorhinal cortex (medial entorhinal cortex, MEC; lateral entorhinal cortex, LEC) and the hippocampus (dentate gyrus, DG; cornu ammonis, CA1, CA2, CA3; dorsal hippocampal commissure, dhc), which includes the subicular complex (subiculum, Sub; pre-/para-subiculum, PPS; post-subiculum, Post).

Our analyses showed that at the early stage, significant volume reduction was observed in the entorhinal cortex of the EC-Tau mice (n = 11, mean age = 20.71 months) compared to their controls (n = 10, mean age = 20.70 months) (Fig. 1a). At the moderate stage, the atrophy in the entorhinal cortex had spread to other regions in the MTL in the EC-Tau mice (n = 10, mean age = 33.38 months) compared to their control group (n = 10, mean age = 32.26 months) (Fig. 1b). More specifically, this significant volume reduction propagated towards the dorsal hippocampal region. Voxel-based analyses showed that atrophy originated from the entorhinal cortex in both the early and moderate stage and spread further into the hippocampus at the later stage. Additionally, we performed voxel-based analyses on EC-Tau mice at a pre-pathological tau stage (6-12 months old, EC-Tau: n = 9, average age = 8.8 months; control: n = 7, average age = 9.2) to test whether tau transgene expression had any effect (Fig. S1). No significant volume reduction was seen throughout the brain.

**Fig. 1.**
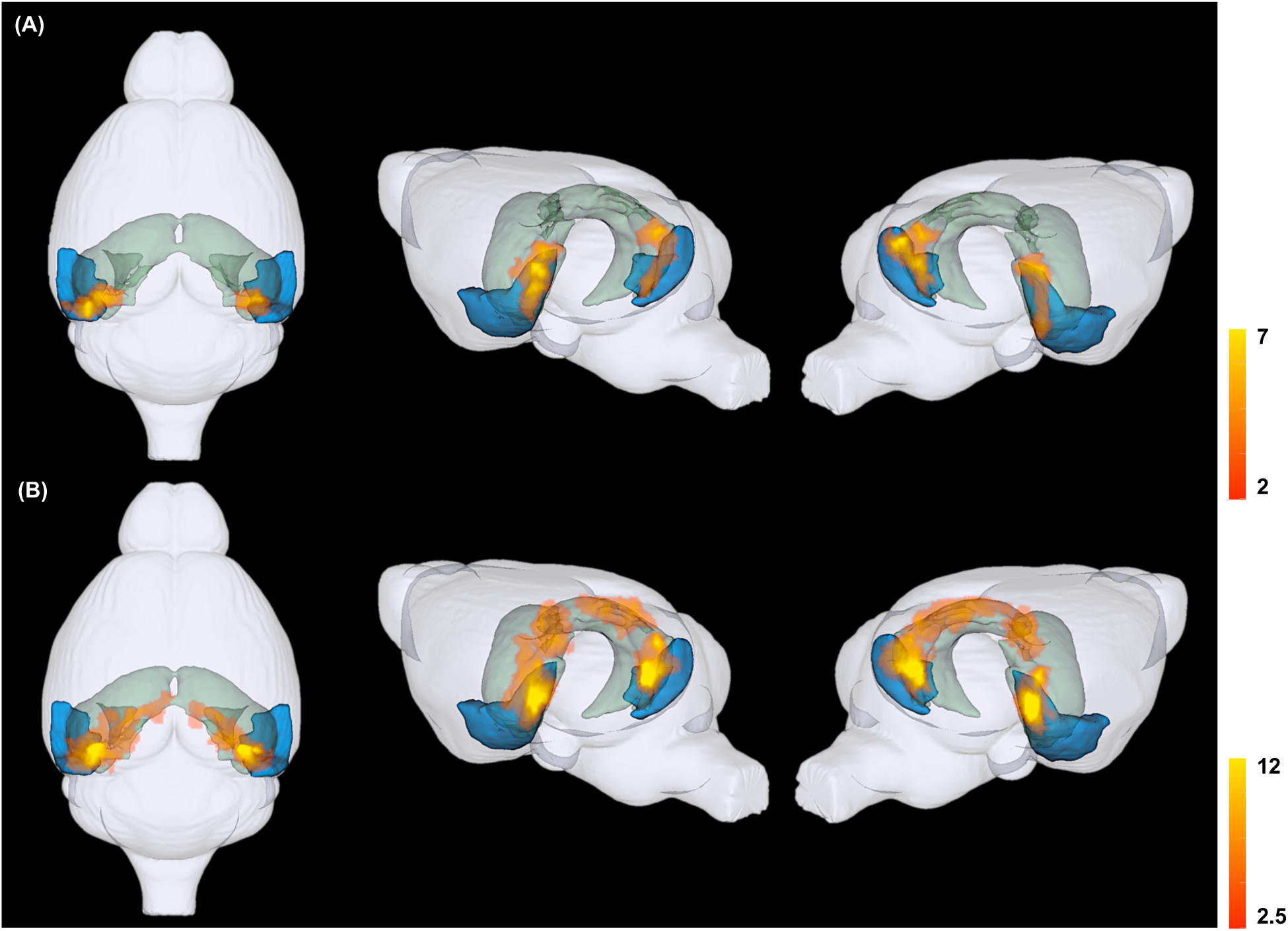
3D rendering of significant volume reduction in EC-Tau mice in the medial temporal lobe (MTL). Tensor-based morphometry (TBM)-related atrophy in the entorhinal cortex (EC) and hippocampus at (**A**) early stage and, (**B**) moderate stage. For each panel, the colored regions depict the EC in blue (LEC, MEC) and the hippocampus (DG, CA1, CA2, CA3, dhc; subicular complex: Sub, PPS, Post) in green. Each panel shows the top view (left), left hemisphere (middle), and right hemisphere (right) of the mouse brain. Voxel-based analyses were conducted using a general linear model in SPM and individual genotypes at each stage were contrasted using two-sample Student’s t test. Statistics are represented as heat maps of *t* values corresponding to voxel-level p < 0.005, cluster-level p < 0.05. Medial entorhinal cortex, MEC; lateral entorhinal cortex, LEC; dentate gyrus, DG; cornu ammonis, CA1, CA2, CA3; dorsal hippocampal commissure, dhc; subiculum, Sub; pre-/para-subiculum, PPS; post-subiculum, Post.

The *in vivo* MRI atlas was also used to identify which specific sub-regions of the MTL were affected. Figure 2 shows two representative 2D axial slices with the MTL atlas labels overlaid on the group-wise template in the top panels (Fig. 2a, 2d), and the significant atrophy in the early and moderate stage in the middle (Fig. 2b, 2e) and bottom panels (Fig. 2c, 2f), respectively. At the early stage, the EC-tau group showed significant volume reduction mainly in the MEC and PPS compared with their age-matched controls. Atrophy spread from these regions and into the hippocampus and subicular complex in the moderate stage.

**Fig. 2:**
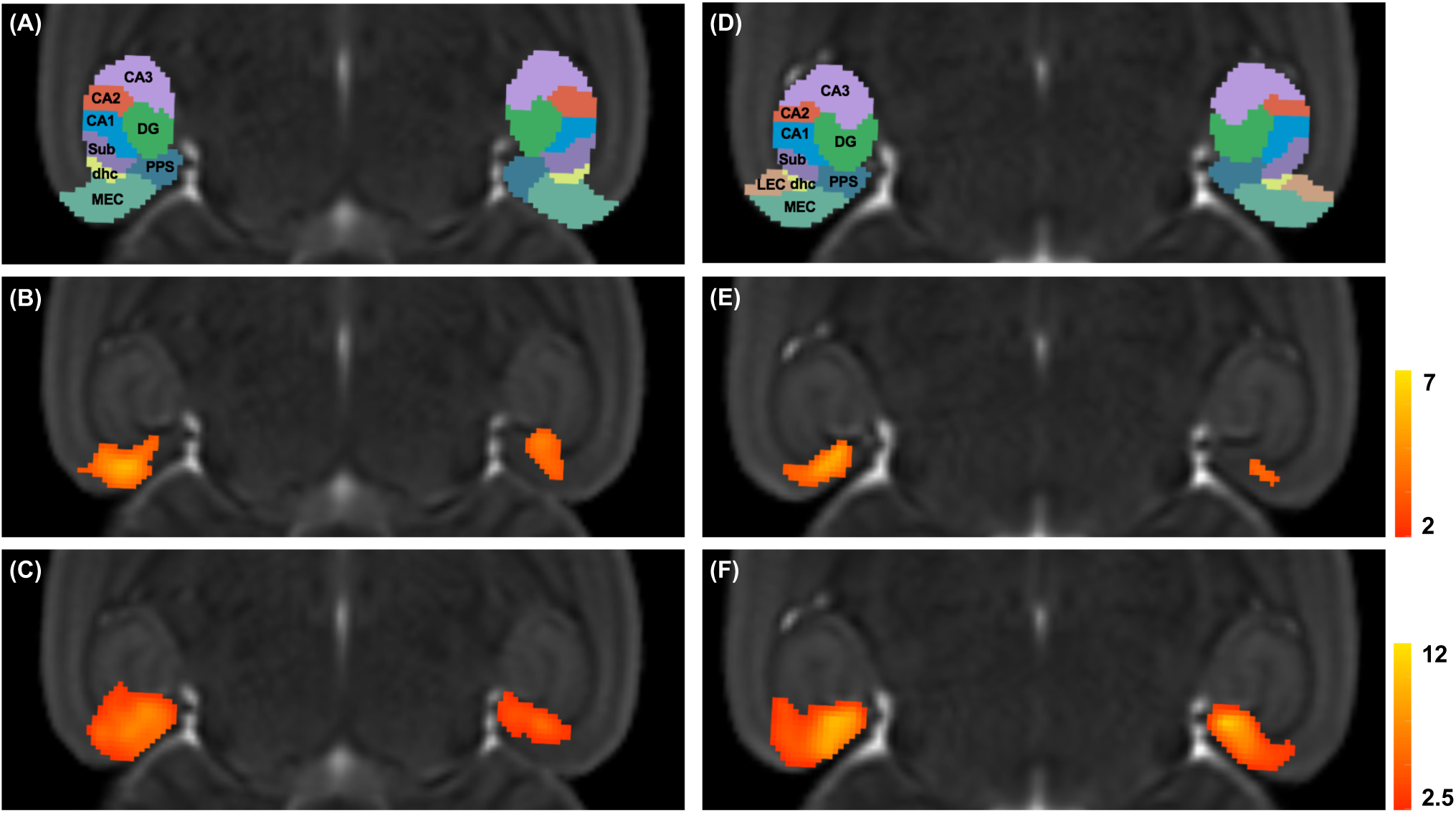
2D representation of significant volume reduction in EC-Tau mice in two axial slices. Tensor-based morphometry (TBM)-related atrophy in a dorsal slice (left) and ventral slice (right). (**A, D**) *In vivo* MRI atlas label overlaid on a group-wise template defining sub-regions of the entorhinal cortex and hippocampus at, (**B, E**) early stage and, (**C, F**) moderate stage. Voxel-based analyses were conducted using a general linear model in SPM and individual genotypes at each stage were contrasted using two-sample Student’s t test. Statistics are represented as heat maps of *t* values corresponding to voxel-level p < 0.005, cluster-level p < 0.05. Medial entorhinal cortex, MEC; lateral entorhinal cortex, LEC; dentate gyrus, DG; cornu ammonis, CA1, CA2, CA3; dorsal hippocampal commissure, dhc; subiculum, Sub; pre-/para-subiculum, PPS; post-subiculum, Post.

### Spread of TBM-related atrophy from EC into hippocampus

To quantify the spread of atrophy in the MTL between the early and moderate stage, we calculated the percentage of each sub-region with significant volume reduction in the 3D volume. Table 1 shows the percentages for each sub-region at both stages. Within the entorhinal cortex, the MEC showed the greatest effect, with 21.52% of the region showing significant volume reduction that increased to 28.72% at the moderate stage. The LEC was minimally affected at both the early and moderate stage. Although tau pathology is apparent in both the LEC and MEC in EC-tau mice, the tau transgene is expressed more abundantly in MEC neurons and there is relatively more intense tau staining in the MEC than in the LEC (*12-14*). Of the hippocampal sub-regions, the CA1, DG, and Sub had the most notable increase in percentage of region with significant atrophy when comparing the two stages. The CA1 volume reduction increased from 0.21% at the early stage to 7.45% at the moderate stage. In the DG, 0.73% of the region was affected at the early stage that increased to 7.65% at the moderate stage. In addition, the subiculum was greatly affected, from 13.45% (early stage) to 44.42% (moderate stage). The PPS, Post, and dhc, which contains projections of pre-subiculum (PrS) from (and to) the EC (*17*), all showed significant volume reduction at the early stage, and the percentage increased at the moderate stage. There was no significant volume reduction in the CA2 and negligible volume reduction in the CA3 at both the early and moderate stage.

**Table 1.** Percentage of each sub-region in the entorhinal cortex and hippocampus with significant volume reduction at the early and moderate stage.

|  | Entorhinal Cortex (%) |  | Hippocampus (%) |  |  |  |  |  |  |  |
| --- | --- | --- | --- | --- | --- | --- | --- | --- | --- | --- |
|  | MEC | LEC | CA1 | CA2 | CA3 | DG | Sub | PPS | Post | dhc |
| **Early** | 21.52 | 0.22 | 0.21 | 0 | 0 | 0.73 | 13.45 | 41.18 | 17.49 | 33.51 |
| **Moderate** | 28.72 | 1.16 | 7.45 | 0 | 0.03 | 7.65 | 44.42 | 55.81 | 22.46 | 50.93 |

### Neuronal loss and volume reduction at the moderate stage

We investigated the relationship between neuronal loss and volume reduction in the EC, PPS, Sub, DG, and CA1 in EC-Tau mice compared to age-matched controls at the moderate stage. Volume reduction analysis was conducted on slices that matched the NeuN stained sections. The number of NeuN^+^ neurons significantly decreased in the EC (t(8) = 4.129, p = 0.003) and in the PPS (t(18) = 6.476, p = 0.0002) in the EC-Tau mice compared to controls, but not in the Sub (t(8) = 0.421, p = 0.685), DG (t(8) = 0.784, p = 0.456), or CA1 (t(8) = 0.034, p = 0.974) (Fig. 3a). There was also significant volume reduction in EC-Tau mice in the EC (t(18) = 3.204, p = 0.005) and in the PPS (t(18) = 7.026, p = 1.479e-6) as well as in the Sub (t(18) = 4.452, p = 0.0003) and the molecular layer of the DG (MoDG, t(18) = 2.628, p = 0.017), but not in the CA1 (t(18) = 0.958, p = 0.351) (Fig. 3b). Thus, the significant volume reduction we observed in the Sub and MoDG precedes overt cell loss at the moderate stage.

**Fig. 3:**
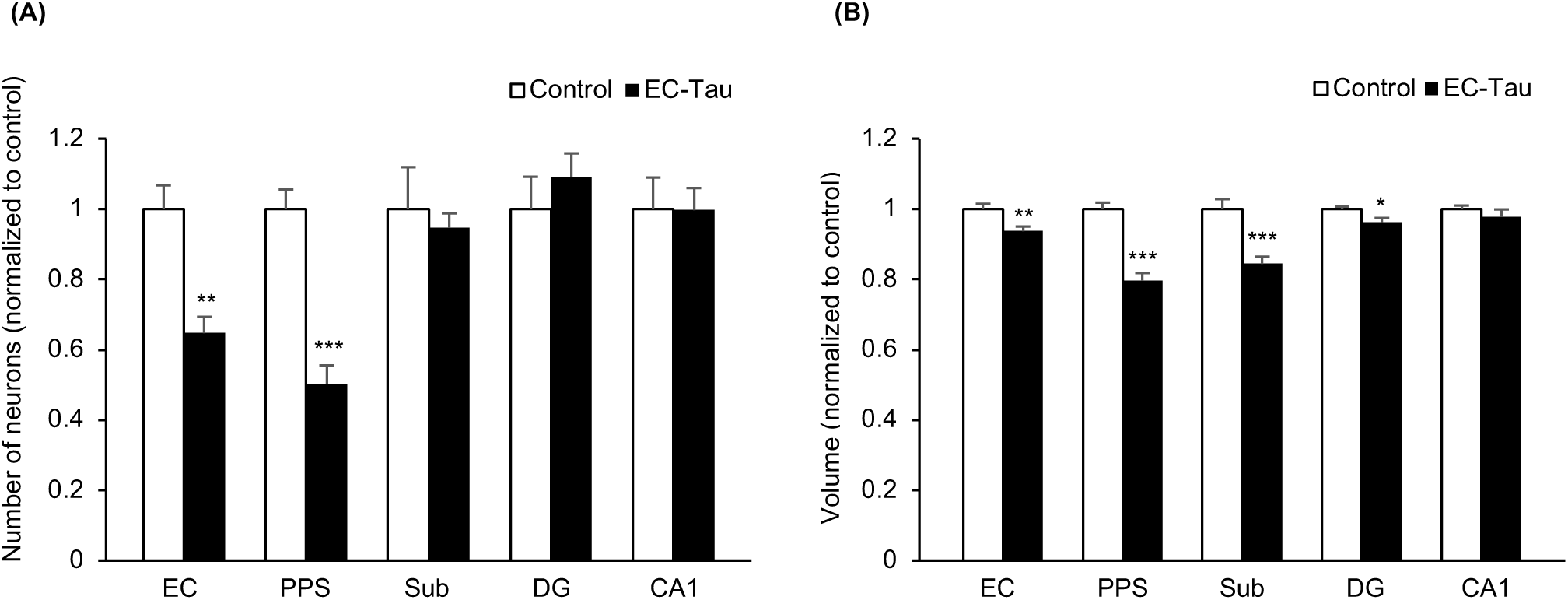
Neuronal loss and volume reduction in EC-Tau mice at the moderate stage. (**A**) The mean number of neurons in each sub-region were compared between control (n = 5) and EC-Tau (n = 5) mice at the moderate stage. (**B**) The mean volume of each sub-region was calculated from the Jacobian determinant and compared between control (n = 10) and EC-Tau (n = 10) mice at the moderate stage. A two-sample two-tailed Student’s t-test was performed to analyze neuron counting and volume reduction. All data are expressed at mean + SEM (*p < 0.05, **p < 0.01, ***p < 0.001). Entorhinal cortex (EC), pre-/para-subiculum (PPS), subiculum (Sub), dentate gryus (DG), cornu ammonis (CA1).

### 3D imaging of iDISCO+ CP27 staining and TBM-related atrophy

To qualitatively visualize the association between tau pathology and significant volume reduction in EC-Tau mice, we co-registered iDISCO+ CP27 (human tau) immunolabeled data from 25-month and 34-month EC-Tau mice with the early and moderate stage atrophy map (Fig. 4). The CP27 antibody detects all human tau, regardless of conformation or phosphorylation status, however, accumulation in the somatodendritic compartment represents pathological tau accumulation. CP27 immunolabeling correlates well with the distribution of tau labeled with the MC1 antibody which detects human tau in an abnormal pathological conformation (*18*). Our results showed human tau immunolabeling throughout the left hemisphere, but the tau labeling was more intense and extensive in the regions corresponding to significant volume reduction in the MTL (Fig. 4a). More specifically, human tau in the MEC and PPS corresponded with the significant atrophy in the same regions at the early stage. Compared to the early stage, less human tau was present throughout the MTL at the moderate stage due to the loss of neurons, but there was significant volume reduction in the EC and PPS and at this stage, in the hippocampus (Fig. 4b).

**Fig. 4:**
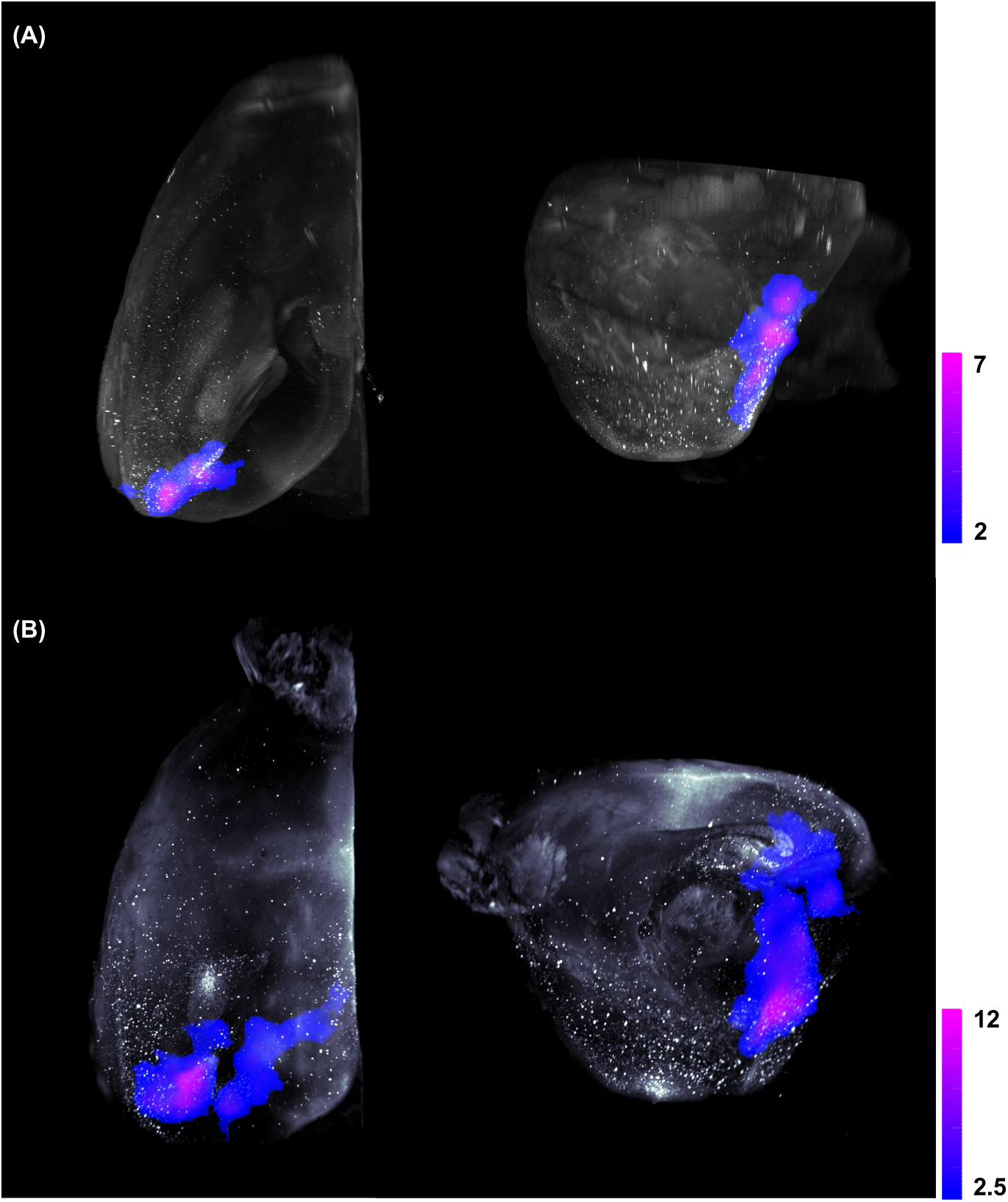
3D representation of iDISCO+ CP27 overlaid with significant volume reduction in the left mouse brain hemisphere. iDISCO+ CP27 (white), which stains all human tau, was co-registered with tensor-based morphometry (TBM)-related atrophy at, (**A**) the early stage and, (**B**) the moderate stage. Each panel shows the top view (left) and left hemisphere (right) of the mouse brain. Voxel-based analyses were conducted using a general linear model in SPM and individual genotypes at each stage were contrasted using two-sample Student’s t test. Statistics are represented as heat maps of *t* values corresponding to voxel-level p < 0.005, cluster-level p < 0.05.

## Discussion

In AD, the first neurons to be affected with tauopathy are in layer II of the EC (*19*). This layer projects directly into the outer two thirds of the molecular layer of the DG (*20*), which together with projections to other subfields of the hippocampal formation form the so-called perforant pathway (*21*). The perforant pathway also consists of projections from neurons in layer III of the EC to the Sub and CA1 (*21*). The Sub and, to a lesser extent, the CA1 form the main output from the hippocampus to the deeper layers of the EC while the para-subiculum (PaS) and PrS also have projections to and from the EC (*22*).

Tensor-based morphometry is a non-invasive, automated imaging technique that has proven to be essential in identifying local structural differences in whole-brain analyses, providing high spatial and temporal resolution to effectively determine the volumetric changes observed at multiple timepoints. In this study, we utilized TBM-MRI to map volume reduction in aging EC-Tau mice at two timepoints that represent two stages of tauopathy in human AD - early and moderate, and to investigate the association of tau pathology with the spread of volumetric reduction and neuronal loss between areas that are synaptically connected to the EC.

Our TBM-MRI data showed that most of the significant atrophy in the EC-tau mouse line was located in the MTL region at both the early and moderate stage, with some affected regions adjacent to the MTL. This could be due to atrophy inside the MTL region causing these adjacent regions to show atrophy as well. By the moderate stage, widespread atrophy projected into the anterior-medial region of the hippocampus and subicular complex. With the whole-brain analysis, we were able to visualize atrophy throughout the entire brain and subsequently determine that the MTL showed the largest region of volume reduction at both the early and moderate stage. Using the co-registered *in vivo* atlas, we identified specific individual sub-regions in the entorhinal cortex and hippocampus with significant atrophy and demonstrated that atrophy originating in the MEC and PPS of the MTL spread into the CA1, DG and Sub regions of the hippocampus and subicular complex.

Delineating the spatial and temporal spread of tau pathology along this pathway has been of significant interest, but the implications for neurodegeneration have not been well explored. *Ex vivo* studies in the EC-Tau mouse line have shown that tau pathology in the EC spreads to functionally connected regions in the hippocampus as the mice age (*12-14*). Age-dependent structural changes such as axonal and synaptic degeneration occur in the entorhinal cortex by 21 months, but with no significant neuronal loss (*12*). By 24 months, both pre- and post-synaptic densities were significantly reduced in the middle third of the molecular layer of the DG (*12*), suggesting synapses are lost in this region as axons originating from neurons in the EC degenerate. In addition, significant neuronal loss was detected in the EC-II and PaS compared to the average neuron number in age-matched control brains (*12*). Axonal tau was dramatically reduced and human tau accumulated in cell bodies in the entorhinal cortex and hippocampus (DG, CA1, CA2/3, Sub, and PrS) at 24 months (*13*). The relocation of tau from the axons to the somatodendritic compartment is one of the earliest events in the pathological cascade of early AD (*2*) and it is possible that axon degeneration is initiated in the EC. The progressive spread of tau pathology associated with the TBM-related atrophy that we see at the early stage (Fig. 1a; Fig. 2b, 2e) suggests that atrophy is caused by axon degeneration and synaptic loss along the perforant pathway from the EC into the DG as well as neuronal loss in the EC and PaS.

EC-tau mice at the moderate stage (approximately 34 months of age) had almost undetectable tau protein in axons but the protein had accumulated in cell bodies throughout the hippocampal formation (*13*). At this age, the number of neurons was significantly reduced in the EC-II and PPS compared to age- and gender-matched control mice, as well as in the EC-III/IV, indicating progressive neuronal loss as pathology worsens and spreads in the brain (*13*). However, there was no significant neuronal loss detected in the CA1, DG, and Sub regions at 34 months in our *ex vivo* study (*13*). Interestingly, our TBM-MRI study showed that the CA1, DG, and Sub regions displayed the greatest increase in percentage of significant volume reduction at the moderate stage (Table 1). We selected slices that matched the NeuN stained sections to further explore the relationship between neuronal loss (Fig. 3a) and volume reduction (Fig. 3b). There was significant neuronal loss and volume reduction in both the EC and PPS. In addition, the Sub and MoDG showed significant volume reduction, but no significant neuronal loss. Our finding indicates that the volume reduction in the Sub and MoDG precedes overt cell loss at the moderate stage.

Neuron loss is associated with the presence of NFTs which are identified as mature tangles by thioflavin S staining (*12-14*). Compared to mice at 24 months of age, larger aggregates composed of thioflavin S positive tau were only observed in the EC and PPS at 34 months (*23*), which correlated with the neuron loss observed in these regions (*13*). This suggests that the volume reduction we observed at the moderate stage in the MoDG and Sub is due to the dysfunction and degeneration of regions directly connected to the EC rather than the presence of NFTs and neuron loss. The atrophy is possibly due to the axon and synapse degeneration initiated in the EC, primarily along the perforant pathway into the molecular layers of the DG. Also, the death of neurons in the EC and PPS could induce degeneration in secondary regions that are directly connected because of the lack of input from the neurons in the EC. In summary, using TBM-MRI we could not only pinpoint specific regions in the brain with atrophy, but we were also able to visualize these changes throughout circuits, in 3D. Importantly, our findings show that not only can we map the structural changes previously observed in *ex vivo* studies, but we can also demonstrate that progressive neurodegeneration precedes overt cell loss in the MoDG and Sub.

Using data generated in our previous study (*13*), we co-registered our 3D TBM-MRI atrophy results with 3D imaging using iDISCO+ immunolabeling of human tau in EC-Tau mice at both the early and moderate stage. Co-registration of human tau pathology and the TBM-related atrophy results mapped the spatial accumulation of human tau to examine the association with structural changes in corresponding regions. Immunolabeled, somatic human tau was most prevalent in the MEC and PPS region that corresponded to the same regions with the highest atrophy in our TBM-MRI results at the early stage. At the moderate stage, less human tau was apparent in the MTL region while there was still an increase and spread of atrophy from the entorhinal cortex into the hippocampus. This finding indicates that degeneration in the hippocampus is subsequent to the degeneration originating in the entorhinal cortex. As discussed earlier, *ex vivo* studies detected neuronal loss in the EC-II and PaS at 24 months (*12*) and in the EC-II, EC-III/IV, and PPS at 34 months, but not in the hippocampal sub-regions such as the CA1, DG, and Sub (*13*). This, together with the co-registered 3D iDISCO+ imaging and TBM-MRI atrophy results further supports the interpretation that the spread of tau pathology is associated with synaptic/axonal degeneration and neuronal loss at the early stage, but at the moderate stage, the atrophy we observe in the MoDG and Sub is not due to neuronal loss. TBM-MRI allowed us to visualize this volume reduction in aged EC-Tau mice, originating from the EC and propagating into the hippocampus.

At both stages, the left hemisphere was more affected compared to the right hemisphere. Several studies have looked at cerebral asymmetry in both rodents and humans. In rodents, significant cerebral asymmetry has been found mainly in the hippocampus, but the asymmetry seems to vary by strain, gender, and age (*24-26*). In human studies, hemispheric asymmetry of the whole hippocampus in healthy adults was also observed, with larger volume in the right hemisphere (*27, 28*). In a study that compared left and right hippocampal volumes in patients with AD and mild cognitive impairment (MCI), the authors showed a consistent asymmetry pattern with a higher percentage of average volume reduction in the left hippocampus of patients with AD and mild cognitive impairment (MCI) compared with aging controls (*29*). The differences between cerebral asymmetry studies could be due to methodological approaches and/or differences in MRI magnet field strength and slice thickness values that might differentially contribute to volumetric asymmetry estimates (*27*). The structural as well as functional laterality differences in cerebral asymmetry in rodents and humans have yet to be defined.

Recent studies in humans examined the association between tau accumulation with PET imaging tracers and cortical atrophy using structural MRI. A significant negative relationship between tracer uptake and concurrent cortical thickness was observed in the MTL (*7-10*), as well as in regions outside of the MTL and in neocortical areas such as the temporoparietal, posterior cingulate/precuneus, and occipital cortices (*3-5, 7-11*). In a recent longitudinal study, results show that the global intensity of tau-PET signal, but not β-amyloid-PET signal, predicted the rate of subsequent atrophy (*11*). In addition, the specific distribution of tau-PET signal was a strong indicator of the topography of future atrophy at the single patient level (*11*). This supports our results showing that the spread of atrophy is associated with tau pathology and demonstrates the utility of transgenic tau mouse models to study the molecular drivers of these effects.

There have been some inconsistencies in the relationship between tau accumulation with PET imaging and hippocampal atrophy with MRI. Some studies have shown that while there was significant atrophy in the hippocampus, there was no significant difference in hippocampal PET tracer retention between the AD patients and controls (*30, 31*). The discrepancy between tau deposits and cortical atrophy may be due to several factors. First, amyloid β (Aβ) status as some studies show atrophy is preferentially associated with tau rather than Aβ pathology (*3-5*). Second, lack of extensive validation of existing tau-specific PET tracers – the type of tau deposits (conformation, maturation stage, tau isoform), their specific binding site(s), and “off-target” binding may affect the sensitivity and specificity of tracers(*32-34*), and third, the absence of longitudinal data in some studies - further longitudinal data is necessary to clarify the spatio-temporal relationships between tau deposition and atrophy (*3, 8, 9, 30*). Our current study overcomes these limitations by using a transgenic tau mouse model to examine the relationship between tau pathology (as opposed to amyloid plus tau pathology), and atrophy, and its association with neurodegeneration, at two stages of disease. It is not possible to separate out these factors in human studies.

In conclusion, we demonstrate that TBM-MRI is an effective, non-invasive imaging technique that can determine which individual regions of the brain are affected at different stages during disease progression. Although the specificity of TBM-MRI does not allow us to identify the exact underlying causes of atrophy, this method is sensitive to detecting atrophy and can guide *ex vivo* experiments in which more conclusive tests can be performed to analyze neuronal vulnerability at a molecular level. Furthermore, while our study focused primarily on volumetric changes in the MTL region at two timepoints, additional research can be conducted to longitudinally track the reduction (or expansion) of volume in other regions with whole-brain analyses. Thus, TBM-MRI can enhance our understanding of the pathological basis and progression of neurodegeneration in Alzheimer’s Disease and other neurodegenerative disorders.

## Materials and Methods

### Study Design

In our study, we conducted a blinded controlled laboratory experiment with transgenic EC-tau mice aged 20-24 months old (early stage) and 30-36 months old (moderate stage) compared with their age-matched littermates. The animal subjects were randomly assigned to experimental groups based on their age (early or moderate stage) and genotype (EC-Tau or control). The animal caretakers provided the mice based on mouse ID number and the animal data was scanned in the MRI and processed randomly. The investigators who assessed, measured, and quantified the results were blinded to the genotype of each group. A power analysis was performed to calculate the necessary sample size for each group. Estimates for group sizes per study were based on effect sizes observed in past experiments investigating volume changes in the brain using the logarithmic transform of the Jacobian determinant. Effect sizes have been approximately 0.08, and standard deviations have been approximately 0.06. These values are used in the following formula to obtain a sample size estimate for an unpaired t-test: N = 1 + 16 x ([standard deviation] / [effect size]) ^ 2. The sample size we arrive at is N = 10 animals for each experimental group and N = 10 for each control group.

The criteria for inclusion and exclusion of data was established prospectively based on SNR (signal-to-noise ratio) of each mouse scan. Typical SNR values range between 15 and 25 (*35*). MRI scans with an SNR in the typical range were included in the data and scans with an SNR outside of this range were excluded from the study. An outlier is a value that is more than three scaled median absolute deviations (MAD) away from the median and was defined before the beginning of this study. No outliers were detected during the study. This study was conducted on a per subject basis and no replicates were applied.

Previous ex vivo studies show structural changes such as axonal and synaptic degeneration in the EC and DG of the hippocampus in EC-Tau mice compared to control mice at the early stage (*12, 13*), while significant neuronal loss was only detected in the EC-II and PaS (*12*). At the moderate stage, tau protein had spread and accumulated throughout the hippocampal formation, but neuronal loss was only seen in the EC and PPS (*13*). Our study aims to delineate the association of tau pathology with the spread of TBM-MRI volume reduction and neuronal loss observed with NeuN+ staining. Furthermore, we qualitatively visualized the association of tau pathology with iDISCO+ CP27 immunolabeled data and our TBM-MRI volume reduction results.

### Transgenic mice

An inducible mouse line in which the expression of human full-length tauP301L (EC-Tau) (*13, 14*) was predominant in the EC was created by crossing the neuropsin-tTA “activator” line (genotype: Tg(Klk8-tTA)SMmay/MullMmmh; strain background: congenic on C57BL/6 background) (*36*) with a tetracycline-inducible “responder” line (genotype: Tg(tetO-MAPT*P301L)#Kha/JlwsJ; strain background: FVB/N background) (*37*) to create the bigenic EC-tau line and control non transgenic littermates (Tg(Klk8-tTA)SMmay/MullMmmh Tg(tetO-MAPT*P301L)#Kha/JlwsJ; strain background: FVB/N:C57BL/6) (*14*). Experimental mice were all F1 progeny. The mice were separated into two age groups representing early (20-24 months old, EC-Tau: n = 11, average age = 20.71 months; control: n = 10, average age = 20.70 months) and moderate (30-36 months old, EC-Tau: n = 10, average age = 33.38 months; control: n = 10, average age = 32.26 months) stages of tauopathy (approximately equivalent to human AD Braak stages I/II and Braak stages III/IV, respectively). Care of transgenic mice was in accordance with protocol approved by the Institutional Animal Care and Use Committee (IACUC) at Columbia University.

### Image acquisition

MRI images were acquired with a 9.4 T vertical Bruker magnet and a 30 mm inner-diameter birdcage radio frequency coil before and 50 minutes after intraperitoneal (IP) injections of contrast agent gadolinium (10 mmol kg^-1^) (*35, 38*). Mice were anesthetized with isoflurane mixed with oxygen delivered through a nose cone (2% for induction, 1-1.5% for maintenance during scan). T2-weighted images were obtained using a fast spin echo sequence with TR = 3500 ms, TE = 15 ms, effective TE = 43.84 ms, in-plane resolution = 86 µm, and slice thickness = 500 µm.

### Image analysis

Using a robust, symmetric method (*39*), an unbiased, within-subject co-registration was performed by iteratively aligning the pre- and post-contrast images to generate a median template image for each mouse (*40*). The median template images were skull-stripped using a pulse-coupled neural network (PCNN) algorithm optimized for rodent brains that operates in 3-D (*41*) to produce whole-brain volumes. The skull-stripped median template images were then upsampled to an isotropic resolution of 86 × 86 × 86 µm^3^ using mri_convert (Freesurfer) with cubic interpolation.

Within-group co-registration was performed using Advanced Normalization Tools (ANTs) (*42, 43*) to transform individual median images into a group-wise template space. First, a linear transformation with 12 degrees of freedom using a cross-correlation (CC) intensity-based similarity metric was optimized to globally align the median images to a randomly selected image in their dataset. Using the linearly aligned images, the optimal Greedy “Symmetric Normalization” (SyN) diffeomorphic transformation was then determined in order to produce the necessary deformations to warp each median image into the group-wise template space. The co-registration algorithm was instantiated by the ANTs script buildparalleltemplate.sh. Finally, the linear transformation and nonlinear warps were applied with the ANTs program WarpImageMultiTransform to transform the median image for each mouse into the group-wise template space. This state of the art SyN method for maximizing CC is a reliable method for normalizing and making anatomical measurements in volumetric MRI in neurodegenerative brain (*43*).

Tensor-based morphometry (TBM) is an image analysis technique that identifies regional structural differences from the gradients of the nonlinear deformation fields, or warps (*15*). The logarithmic transformation of a Jacobian field is a common metric used to evaluate these structural differences at a voxel level and has become a standard in TBM (*44-46*). Using the ANTs program ANTSJacobian, the Jacobian determinant (JD) of the warp and the logarithmic transform of the JD of the warp for each mouse was generated in order to determine the structural changes between the EC-Tau mice and controls for each age group.

### Atlas-based segmentation

An *in vivo* MRI atlas of the mouse brain was constructed to determine which specific sub-regions of the brain are affected. To do this, an *ex vivo* MRI mouse brain atlas (Australian Mouse Brain Mapping Consortium, AMBMC) with hippocampus (*17*) and cortex (*47*) labels was utilized. Sub-regional segmentation was fitted to the symmetric model (*48*) to avoid left/right average differences. First, the symmetric model mouse brain, hippocampus labels, and cortex labels were downsampled to an isotropic resolution of 86 x 86 x 86 µm^3^ using mri_convert (FreeSurfer). Similarly, the group-wise template was upsampled to a matching isotropic resolution. The hippocampus and cortex regions were manually masked on both the symmetric model mouse brain and group-wise template to yield volumes containing regions of interest. Using the ANTs script mentioned earlier, the masked symmetric model brain was registered to the masked group-wise template using a linear transformation with a CC similarity metric, followed by a Greedy SyN transformation to determine the necessary deformations to warp the symmetric model brain into the group-wise template space. With the ANTs program, the linear transformation and nonlinear warps were applied to the hippocampus and cortex labels and overlaid on the group-wise template image in 3DSlicer (http://www.slicer.org).

### Voxel-based statistical analysis

Voxel-based analyses were conducted using a general linear model in Statistical Parametric Mapping (SPM, Wellcome Trust Centre for Neuroimaging). The logarithmic transform of the JD maps of individual genotypes at each stage (early: n = 11 EC-Tau, n = 10 control; moderate: n = 10 EC-Tau, n = 10 control) were contrasted using two-sample Student’s *t* test. Results were corrected for multiple comparisons using Monte Carlo simulation implemented in AFNI-AlphaSim (https://afni.nimh.nih.gov/afni/) with 5,000 iterations to achieve a voxel-wise p < 0.005 and cluster-wise p < 0.05. The thresholded *t* maps from individual group comparisons were then overlaid on the group-wise template image in 3DSlicer (http://www.slicer.org) with the registered hippocampus and cortex labels.

### Percentage of sub-regions with significant volume reduction

The percentage of each sub-region with significant volume reduction in the entorhinal cortex and hippocampus were calculated to quantify the spread between the early and moderate stage. First, we used the thresholded *t* map for each age group to create a binary image labeled with value 1 for voxels with significant volume reduction and value 0 for all other voxels that were not affected. Using the *in vivo* MRI atlas and a custom MATLAB script, we calculated the number of voxels in each sub-region of the MTL labeled with value 1 and divided by the total number of voxels in each sub-region to determine the percentage of voxels affected in both the early and moderate stage.

### Neuronal counts and volume reduction analysis at moderate stage

Data from neuronal counts was modified from Fu, et. al., 2016 and incorporated in our study. Briefly, mouse brains (n = 5 controls, n = 5 EC-Tau mice at ∼34 months old) were harvested and drop-fixed in 4% PFA at 4°C overnight, followed by incubation in 30% sucrose (Sigma-Aldrich, Saint Louis, MO, USA). OCT-embedded brains were sectioned (35 µm) throughout on a horizontal (axial) plane with a cryostat (Leica CM3050S, Leica Biosystems, Buffalo Grove, IL, USA), and collected in individual wells. Every 9th free-floating section was selected and stained with mouse anti-NeuN antibody (EMD Millipore, Billerica, MA, USA; 1:1000), which is a neuronal marker. A semiquantitative count of NeuN+ neurons in the EC, PPS, Sub, CA1, and DG was performed in the above selected sections. For each mouse, a total of 10 NeuN stained horizontal sections starting from Bregma -2.04 mm, spaced at 300 µm, were included for automated cell counting (http://imagej.net/Particle_Analysis) using the ImageJ software (version 1.48, US National Institutes of Health, Bethesda, Maryland, USA).

Similarly, the co-registered JD of the warp for each mouse in the moderate stage (n = 10 control, n = 10 EC-Tau, 30-36 months old) were used to observe the relationship between atrophy and NeuN+ neurons in the same regions as mentioned above. Slices from the JD of each mouse were selected to match the NeuN stained sections. Using a custom MATLAB code and the MRI atlas, the sum of the JD for each sub-region was calculated for each mouse in the moderate stage.

### Statistical analysis

The data is expressed as the mean +/- the standard error of the mean (SEM). A two-sample two-tailed Student’s *t*-test was used to analyze the neuronal counting (n = 5 controls, n = 5 EC-Tau mice) and the volume reduction in mice (n = 10 controls, n = 10 EC-tau) at the moderate stage. A value of p < 0.05 was considered statistically significant.

### iDISCO+ CP27 staining and TBM-related atrophy co-registration

As previously described (*13*), the 3D iDISCO+ immunolabeled datasets of Alexa Fluor 647-labeled CP27 antibody (conformation and phosphorylation status independent) in a 25 month EC-Tau mouse and a 34 month old EC-Tau mouse were used to represent the early stage and moderate stage, respectively. Using the Imaris software, the 3D iDISCO+ datasets were exported as a stack of 2D slices. The datasets were downsampled to an isotropic resolution of 86 x 86 x 86 µm^3^ using mri_convert (Freesurfer) to match the resolution of the MRI group-wise template. Co-registration between the iDISCO+ dataset and their respective age-matched group-wise template was performed using Advanced Normalization Tools (ANTs) as mentioned previously. A linear transformation with 12 degrees of freedom using a CC similarity metric was implemented to align the two datasets. Using the linearly aligned datasets, the optimal Greedy SyN diffeomorphic transformation was then determined in order to produce the necessary deformations to warp the group-wise template into the iDISCO+ data template space. Finally, the linear transformation and nonlinear warps were applied to the voxel-based thresholded t maps for both age groups. This transformation co-registered the voxel-based atrophy results with their age-matched iDISCO+ pathology data. The co-registered atrophy data was overlaid on the iDISCO+ pathology data and visualized in 3D using the Imaris software.

## List of Supplementary Materials

Fig. S1. 2D representation in axial view of significant volume reduction at a pre-pathology stage in EC-Tau mice compared to age-matched controls.

## Acknowledgements

We acknowledge Dr. H. Fu (Ohio State University) for assistance with iDISCO+ immunolabeling data. Dr. Scott Small and Dr. Usman Khan (Columbia University) are thanked for their contributions to some of the data used in these studies.

## Author Contributions

This study was designed and managed by C.F., J.G., E.K., and K.D. Animal care and breeding was performed by H.F. Mouse MRI acquisition and analysis was performed by C.F. and J.G. Manuscript preparation was performed by C.F., with input from C.F., J.G., E.K., and K.D.

## Data and Materials Availability

The datasets generated during and/or analyzed during the current study are available from the corresponding author.

