## Supplementary Figure 1 for "Atrophy associated with tau pathology precedes overt cell death in a mouse model of progressive tauopathy"

**Supplementary Materials**

**Fig. S1**

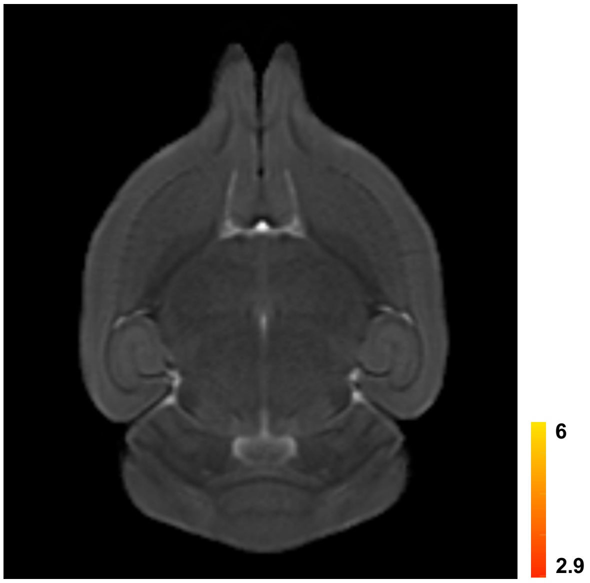


**Fig. S1:** 2D representation in axial view of significant volume reduction at a pre-pathology stage in EC-Tau mice compared to age-matched controls. No significant volume reduction was observed throughout the brain at this stage. Voxel-based analyses were conducted using a general linear model in SPM and individual genotypes at each stage were contrasted using two-sample Student's t test. Statistics are represented as heat maps of *t* values corresponding to voxel-level $p < 0.005$, cluster-level $p < 0.05$.
